# Coalescent times and patterns of genetic diversity in species with facultative sex: effects of gene conversion, population structure and heterogeneity

**DOI:** 10.1101/019158

**Authors:** Matthew Hartfield, Stephen I. Wright, Aneil F. Agrawal

## Abstract

Many diploid organisms undergo facultative sexual reproduction. However, little is currently known concerning the distribution of neutral genetic variation amongst facultative sexuals except in very simple cases. Understanding this distribution is important when making inferences about rates of sexual reproduction, effective population size and demographic history. Here, we extend coalescent theory in diploids with facultative sex to consider gene conversion, selfing, population subdivision, and temporal and spatial heterogeneity in rates of sex. In addition to analytical results for two-sample coalescent times, we outline a coalescent algorithm that accommodates the complexities arising from partial sex; this algorithm can be used to generate multi-sample coalescent distributions. A key result is that when sex is rare, gene conversion becomes a significant force in reducing diversity within individuals, which can remove genomic signatures of infrequent sex (the ‘Meselson Effect’) or entirely reverse the predictions. Our models offer improved methods for assessing the null model (I.e. neutrality) of patterns of molecular variation in facultative sexuals.

## Introduction

Facultatively sexual species, which have both parthenogenetic and sexual stages in their life-cycles, are widespread in nature. They have been a focus for empirical studies of the role of sex in evolution (Goddard *et al.* 2005; D’SOUZA and Michiels 2008; KING *et al.* 2009; Morran *et al.* 2009; Becks and Agrawal 2010; Hojsgaard and HÖrandl 2015). The biology of facultative sexuals is also a research field with broad applications, since many organisms of evolutionary, medical and agricultural importance undergo both sexual and asexual reproduction (Chang *et al.* 2013; Ankarklev *et al.* 2014; Hand and Koltunow 2014; Yoshida *et al.* 2014). However, our understanding of structuring of genetic diversity in such organisms remains limited.

For effective population genetic analysis of facultative sexual individuals to be made, there needs to be a theoretical basis for predicting how neutral diversity is affected under various demographic and reproductive scenarios. This is so that the two are not confounded, especially since they are strongly intertwined (reviewed by Halkett *et al.* (2005); Arnaud-Haond *et al.* (2007)). A classic prediction for organisms with very low rates of sex is that, due to the resulting lack of segregation, diploid asexuals tend to accumulate extensive diversity within individuals, since the two alleles from a single individual remain isolated by descent in the absence of sex. This phenomenon has been labelled the ‘Meselson’ effect, since it was used by Meselson and colleagues to demonstrate a lack of sex in bdelloid rotifers (Mark Welch and Meselson (2000); Butlin (2002); but see Mark Welch *et al.* (2008) for an alternative explanation for their original findings).

One theoretical approach to quantify this effect has been to derive how traditional diversity measures, such as the effective population size *N*_*e*_ and *F* statistics, are affected when sex is rare (Balloux *et al.* 2003; Yonezawa *et al.* 2004; de Meeûs and Balloux 2005). Such studies are based on pairwise sample comparisons, with heightened within-individual diversity not becoming apparent unless sex is extremely rare (with frequency less than 1*/N*, for *N* the total population size). Combining these summary statistics with information on clonal or genotypic diversity can improve inference when rates of sex are not too low (Balloux *et al.* 2003; Halkett *et al.* 2005; Arnaud-Haond *et al.* 2007).

An alternative approach is to analyse polymorphisms using coalescent theory, to recreate the possible genealogies of samples (Kingman 1982; Wakeley 2009). Basic calculations were carried out by Brookfield (1992) and Burt *et al.* (1996) to determine how partial sex affects diversity in DNA fingerprinting analyses and isolates of a human pathogen respectively. More complete analyses were then performed by Bengtsson (2003) and Ceplitis (2003) to determine coalescence times for pairs of samples taken either from the same or different individuals. These analyses confirmed that the rate of sex has to be less than the reciprocal of the population size for coalescent times to be substantially altered, compared to the classic case of obligate sex.

With the advent of high-throughput genome sequencing technologies, there are growing opportunities to characterise both the demographic and reproductive history of partially sexual populations. However, formal theory does not yet exist for genealogies of facultative sexuals that take into account demographic factors, gene conversion, and temporal and spatial heterogeneity in the rate of sexual reproduction. The potential importance of gene conversion is especially pertinent, since there is growing recognition that this process may be an important homo-genising force in diploid asexual populations (Crease and Lynch 1991; SCHÖN *et al.* 1998; Normark 1999; Schön and Martens 2003; Schaefer *et al.* 2006; Flot *et al.* 2013). In addition, previous theoretical studies have not accounted for cases where the rates of sex change in either space or time. It is well known that several organisms adjust their rate of sex in a stressful or dense environment. This mechanism is well-supported both theoretically (Redfield 1988; Hadany and Otto 2007) and empirically (Grishkan *et al.* 2003; Snell *et al.* 2006; Levin and King 2013), but the overall consequences for patterns of genetic diversity are not clear. Previously published theory also suffers from containing ambiguities or typographical errors (explained below), which make interpretation of existing results difficult. In addition, no algorithm yet exists for how to recreate genealogies for multiple samples with partial asexuality, which is essential for simulating the distribution of neutral variation under complex scenarios.

Here, we outline theory to rectify this issue. We rederive equations of the coalescent time for pairs of samples taken either from different or the same individuals, using a structured coalescent framework (Nordborg 1997), clarifying discrepancies that appear between Bengtsson (2003) and Ceplitis (2003). We further show how additional phenomena, including self-fertilisation, gene conversion, population subdivision, and heterogeneity in rates of sex, can be included into the coalescent with partial sex. These results can be used to derive estimators for mutation, migration, and sex rates, based on mean pairwise diversity. These initial analyses will then inform on how to create a simulation program where there exists an arbitrary rate of sex, which can be used to efficiently create genealogies for multiple samples.

## The Basic Model

We begin by considering a population with an arbitrary rate of sex, but is otherwise ideal. Bengtsson (2003) and Ceplitis (2003) derived coalescent times for two alleles (“two-sample” coalescent times) from this basic model. However, we reexamine this model to clarify discrepancies between the earlier studies, and to also lay the groundwork for further extensions.

### Model outline

We consider a diploid Wright-Fisher population of size *N* individuals, so there are 2*N* gene copies in total. *σ* is the rate of sex; that is, with probability *σ*, an offspring is produced sexually, otherwise it is produced asexually. For a specific individual, one can imagine that each of its gene copies inhabit separate ‘genomic demes’ within an individual, pertaining to the left-and right-hand allele copy respectively (Ceplitis 2003). We choose two of these gene copies, and track their genealogy back in time until they reach their common ancestor. These allele pairs can either be sampled from different individuals, or the same individual. With obligate sex this distinction is irrelevant, due to the random segregation and union of gametes. However, if sex is infrequent, then pairs of alleles can remain ‘packaged’ within an individual lineage as a left and right haplotype for long periods of time. This structuring, and the low level of exchange between haplotypes when sex is low, creates a fundamental difference from the standard coalescent.

Two sampled alleles can be found in one of three states: (i) one allele in each of two separate individuals; (ii) two separate alleles within the same individual; and (iii) coalesced. Ceplitis (2003) argued that it is necessary to divide the first state into two separate states: both alleles in the same deme (either the ‘left’ or ‘right’ demes of separate individuals), or one in left and one in right. However, it can be shown that this partitioning is unnecessary as long as the appropriate transition probabilities are applied, which average over these possible states. We calculate the coalescent time by first determining the transition probabilities among the three states mentioned above. We assume that *N* is large, so terms of order 1*/N*^2^ and higher are discarded.

The transition matrix for the process is:

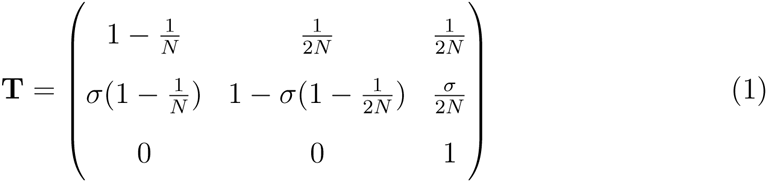

Further details of all transitions and analysis are outlined in Section S5 of the Supplementary Derivations file.

To find the expected coalescence times, we follow the method of Slatkin (1991). Note that the 2 *×* 2 sub-matrix **G**, representing the top-left corner of **T** (Equation 1) is a matrix of non-coalescence; that is, it denotes the probability that two samples do not coalesce in a single generation. The probability that two alleles from either state have not coalesced *t* generations in the past is given by **a**(*t*) = **G**^*t*^_**a**_(0), where **a**(0) = [1, 1]^*T*^.

Conversely, the probability that two alleles *have* coalesced by time *t* is *P* (*t*) = **a**(*t*−1)−**a**(*t*), which can be used to calculate the complete distribution of coalescent times. However, here we will focus on the mean coalescent time; Slatkin (1991) showed that it can be calculated by:

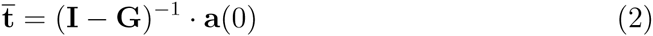

where **I** is the identity matrix. The non-coalescence matrix **G** is crucial for determining the expected coalescent times. It can be decomposed into the relative contributions of sexual and asexual reproduction as follows:

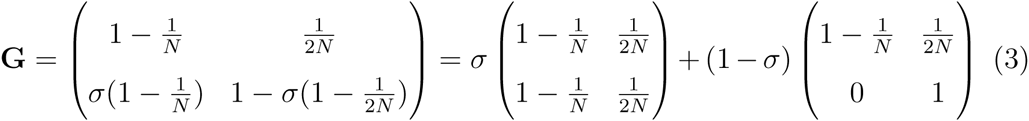

This decomposition makes it easier to see how other complexities (e.g. selfing, gene conversion) can be included in this framework.

Using Equations 2 and 3 we obtain the coalescence times for two samples taken from different individuals, *t*_*b*_, and from the same individual, *t*_*w*_:

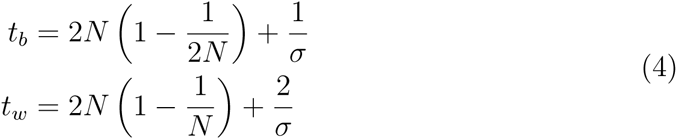

Ceplitis (2003, Equation 2) derived equivalent equations for the limiting case of *σ* = *O*(1*/N*); Bengtsson (2003) presented results in terms of the eigenvalues of **G** (his Equations 1 and 2), although there appears to be typographical errors in the way these eigenvalues are presented. In Section S1 of the Supplementary Mathematica file, we show how equivalent results can be derived using the eigenvalues of **G**, and how they compare to what is presented in Bengtsson (2003).

Visual inspection of Equations 4 shows some key properties of the within-and between-individual coalescent times. It is clear that the rate of sex will only significantly affect the coalescence time if it is very rare, at least *O*(1*/N*), as found in previous studies. Secondly, the within-individual coalescence time is greater than that between-individuals, formalising the ‘Meselson effect’ (Mark Welch and Meselson 2000; Butlin 2002). Two samples from within a single individual are, by definition, members of different ‘left/right’ sides. Sex is therefore required to put them in the same side before coalescence can occur (on average two sex events are needed). Only 50% of two-allele samples from different individuals will be in different ‘left/right’ sides, so the increase in mean coalescence time due to low rates of sex for between individual samples is only half as large.

Figure 1(a) demonstrates how the coalescent times rapidly increase as *σ* becomes significantly less that 1*/N*. If we set *σ* = 1*/N* into Equation 4, then *t*_*b*_ = 3*N -* 1 and *t*_*w*_ = 4*N -* 2. That is, the between-individual coalescent times are around 1.5 times higher than the Kingman (obligate sexual) coalescent, the within-individual coalescent times are twofold higher.

**Figure 1:**
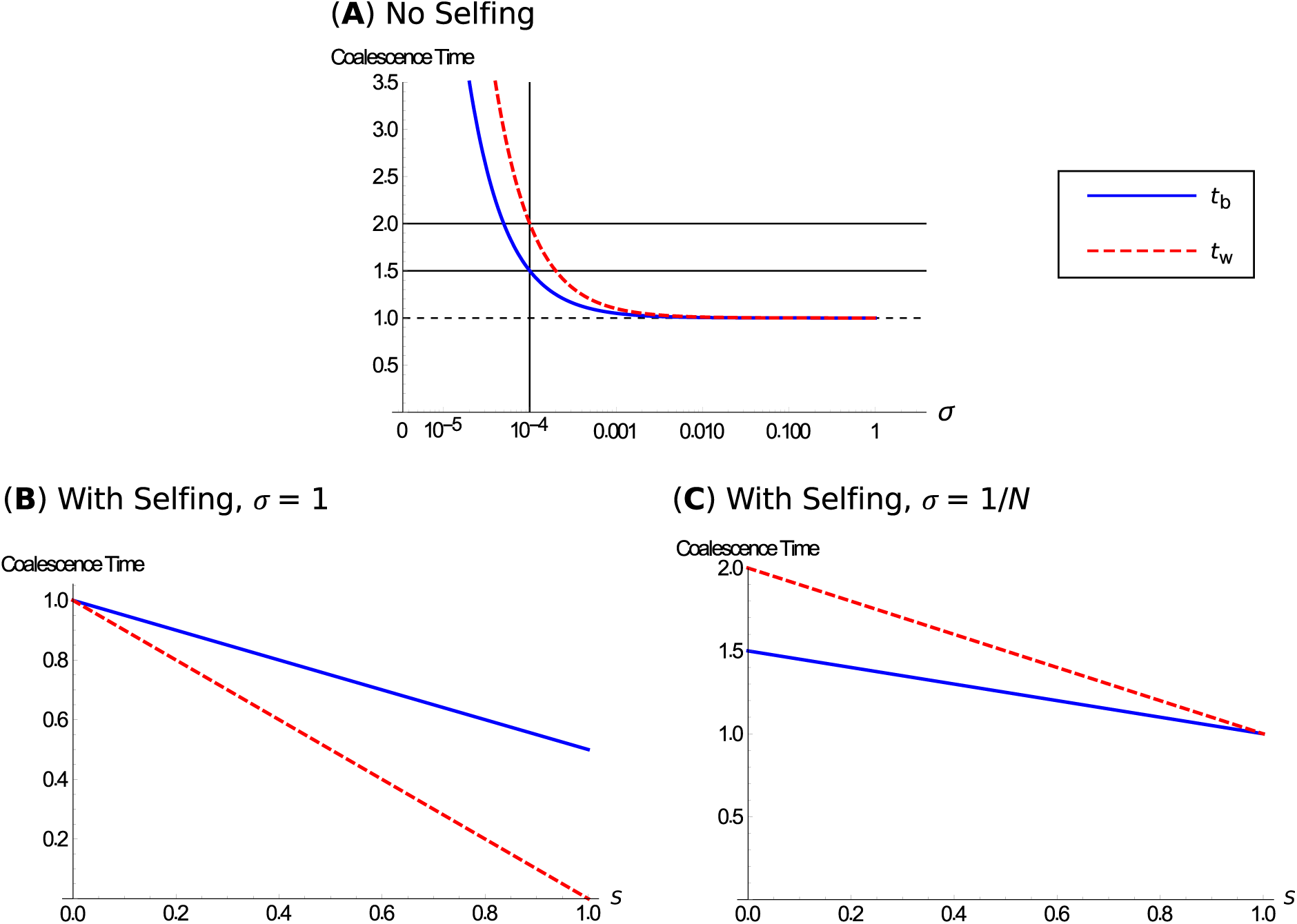
Coalescence times with different reproductive systems. The population size for all plots is *N* = 10, 000. (a) plots coalescence time according to Equation 4, scaled to 2*N* generations, as a function of sex, *σ*. The blue line represents between-individual coalescence time *t*_*b*_, the red line is within-individual coalescence time *t*_*w*_. The vertical line represents *σ* = 1*/N*; the horizontal solid lines show the predicted coalescence times for this *σ* value (*t*_*b*_ *≈* 3*N*; *t*_*w*_ *≈* 4*N*), while the horizontal dashed line shows *t* = 2*N*. (b), (c) show coalescence time if selfing is also included, according to Equation 6. Line colours are the same as in (a), with *σ* = 1 (b) or *σ* = 1*/N* (c).

## Multisample coalescent simulations

The two-sample results are useful at clarifying how neutral drift should operate in facultative sexual populations under a variety of scenarios. However, when analysing empirical data, one would normally obtain multiple samples and then estimate neutral diversity based on these. For obligate sexual organisms, coalescent simulations have proved essential at modelling expected neutral diversity levels as caused by complex demographic scenarios, such as population bottlenecks or expansions (Hudson 2002). Equivalent analysis on sequence data from facultative sexual species is not possible using existing coalescent simulation software. This is because gene samples remain paired within individual lineages if sex is rare; a type of genetic isolation not considered in existing simulation packages. Therefore, it is necessary to create a new coalescent algorithm to account for partial rates of sex.

In Section S6 of the Supplementary Derivations file, we describe how to create a coalescent algorithm that includes partial rates of sex. Standard coalescent algorithms generally proceed in two stages (Wakeley 2009): first, the expected time until an event (state change) is drawn from an exponential distribution, the mean of which is dependent on the total probability that any state change arise, including coalescence. Given this time, the actual change that occurs is subsequently determined by the conditional probabilities of each possible state change, given that a change arises. A third step can be added in which mutations are added to the resulting genealogy in proportion to the branch lengths.

To allow for any rate of sex, we modify the procedure as follows. One draws the expected time until either a state change involving at least one sex event occurs, or a state change not involving sex. Once this time is drawn, it is then determined if the change involved sex or not; if so, the number of sexual splits is drawn from a zero-truncated binomial distribution. The number of trials is determined by the current number of paired within-individual samples, and the probability of sex *σ*. After sexual reproduction is resolved, it is then specified what, if any, other events occur that generation. Hence unlike in standard coalescent simulations, there can be multiple state changes, especially if rates of sex are not too low. The process is repeated until all samples coalesce. We have implemented this algorithm into a simulation program for R (R Development Core Team 2014), which is available as Supplementary Material S8 or can be downloaded online (*URL to be confirmed*).

Using this simulation, we investigated how traditional population genetic parameters are affected under low rates of sex, to determine if there is a risk that they show spurious signs of selection or demography. We ran our coalescent algorithm while varying the rate of sex, and analysed the mutational outputs to calculate summary statistics. 1000 trees were ran for each point for all simulation results throughout. Confidence intervals were calculated using a normal-distribution approximation; similar intervals were produced if using 1000 bootstrap samples.

Results are shown in Figure 2 for *N* = 10, 000, *θ* = 5 with 25 paired samples, although results are nearly identical if 50 single samples were simulated instead (Figure S1 of the Supplementary Derivations file). As the rate of sex decreases and drops below 1*/N*, the number of segregating sites shoots up due to the increased coalescent times, especially within individuals, creating greater diversity (Figure 2(a)). This leads to an increase in traditional estimates of *θ*, which are much higher than the true mutation rate (Figure 2(b)). With rare sex, Tajima’s D also increases to high values (Figure 2(c)), while Fay and Wu’s H drops despite a small increase for *σ ≈* 1*/N* (Figure 2(d)). Both these behaviours are caused by a increase in the number of intermediate-frequency variants, especially within individuals, as sex becomes rare. Finally, the number of unique haplotypes increases (Figure 2(e)), and the number of unique genotypes decreases (Figure 2(f)) as *σ* drops. Both plots reflect how the lack of segregation creates new haplotypes within individuals, but homogenises individual genotypes, in line with previously-reported theory by Balloux *et al.* (2003). These results make clear that if one does not account for rare rates of sex, spurious signatures of selection or demography can arise that can confound analyses. Deeper branches can also lead to an over-estimation of the mutation rate or effective population size.

**Figure 2:**
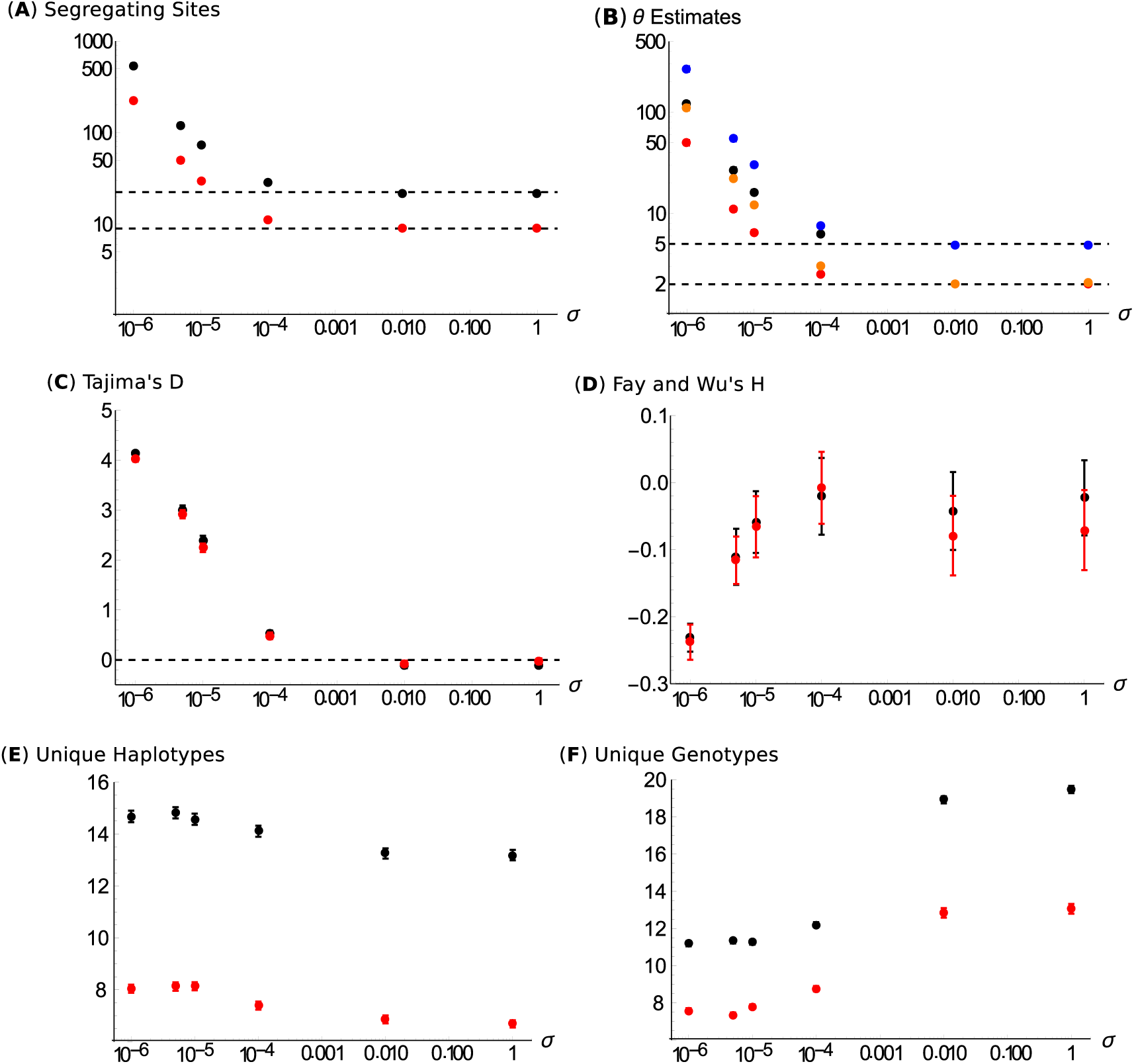
How classic genomic summary statistics are affected by infrequent sex in a panmictic population. All graphs are as a function of *σ*; parameters are *N* = 10, 000, *θ* = 5 (black points by default) or *θ* = 2 (red points by default), based on 25 paired samples, averaged over 1000 coalescent simulations. Error bars are 95% confidence intervals; if they cannot be seen, they lie within the points. (a) Mean number of segregating sites in a sample; the horizontal bar shows the expectation for *σ* = 1 (obligate sex). (b) Estimates of *θ* using Watterson’s *θ* (black or red points for *θ* = 5 or 2), or the mean pairwise differences between pairs of samples (blue or orange points for *θ* = 5 or 2). (c) Tajima’s D. The horizontal line at zero is the unbiased expectation for obligate sex. (d) Fay and Wu’s normalised H statistic. (e) The mean number of unique haplotypes present. (f) The mean number of unique genotypes present.

### Sex with selfing

We can extend the above analysis to consider a case where, if an individual reproduces sexually, it can self-fertilise with probability *S*, and outcross with probability 1 *-S*. The coalescent for a strictly outcrossing-selfing case has been well-investigated (Milligan 1996; Nordborg and Donnelly 1997; Nordborg 1997, 2000; Nordborg and Krone 2002); the key result is that the genealogy is equivalent to a Kingman coalescent, but with the population size scaled by the selfing rate *N*_*e*_ = *N* (2 *-S*)*/S*, so coalescence time is reduced. We can implement similar methods to determine how inbreeding and asexuality interact to affect coalescence times.

The non-coalescence matrix (**G**) with selfing and asexuality is written as (see Section S5 of the Supplementary Derivations file for further information):

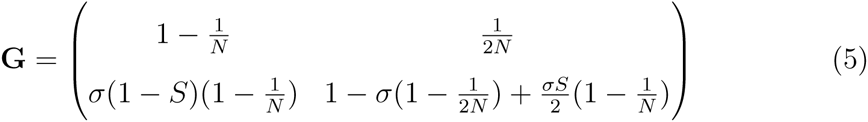

Using Slatkin’S (1991) matrix-inversion method, the coalescent times are obtained as:

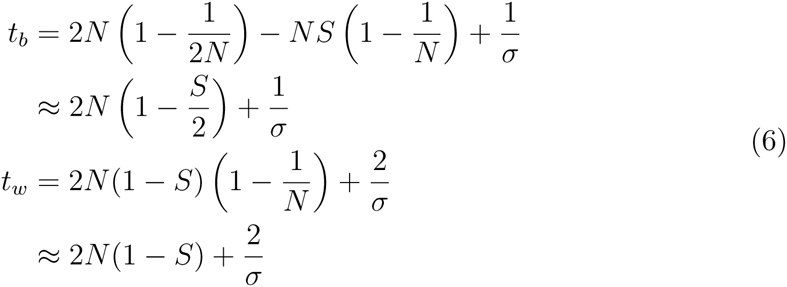

Approximate solutions arise in the limit of large *N*. Equation 6 makes explicit how selfing reduces the coalescent times. Ceplitis (2003, Equation 5) derived equivalent equations in the limit of low rates of sex. However, there appears to be a typographical error in his term for coalescence for two samples, taken from different individuals and different chromosome arms. Similarly, Bengtsson (2003) derived accurate terms for the eigenvalues of **G** in the limit of weak sex, but plots of the effect of selfing appears to be inaccurate (compare his Figure 2 with our Figure 1(c)). These differences are discussed further in Section S1 of the Supplementary Mathematica file.

Selfing and asexual reproduction have opposing effects on the coalescent times, and differently affect *t*_*b*_ and *t*_*w*_. With selfing, *t*_*b*_ > *t*_*w*_ when rates of sex are high, but the reverse is true when the rates of sex are low (Fig. 1b and 1c). With no asexual reproduction, complete selfing reduces the between-individual coalescent time by half, while the within-individual coalescent time is instantaneous (Figure 1(b)). Similarly, for *σ* = 1*/N*, selfing can reduce both sets of coalescent times so that they equate to 2*N* with complete selfing (Figure 1(c)).

### Gene conversion

Gene conversion can strongly affect the genetic architecture of organisms, and can be especially important in facultative sexual organisms, as recently shown in the genome sequence of bdelloid rotifers (Flot *et al.* 2013). If a gene conversion event arises where one allele sample replaces another during reproduction (irrespective of reproduction type), only one of the two chromosome arms is passed on to its offspring. We assume unbiased gene conversion occurs at rate *γ*. The non-coalescence matrix becomes:

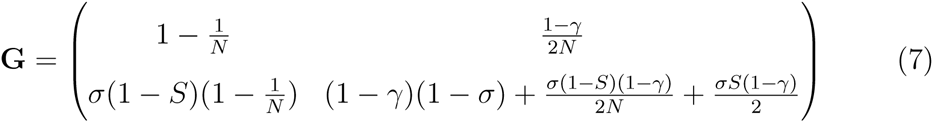

We can solve this matrix as before, and find that:

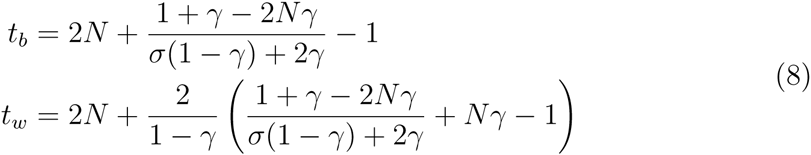

Solutions with selfing are provided in Section S1 of the Supplementary Mathematica file. Assuming gene conversion and rates of sex are small but not too small relative to population size (i.e. 1*/N ≪ σ*, *γ ≪* 1), then Equation 8 becomes:

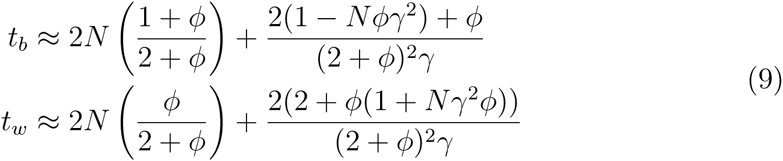

where *ϕ* = *σ/γ*. Under the specified assumptions, the first term in each equation is expected to dominate the expression. If sex is rare relative to gene conversion (*ϕ →* 0), these leading terms go to *N* and 0, respectively; for *t*_*w*_ we must then consider the second term, which goes to 1*/γ* as the first term goes to zero. This shows that low sex with comparatively high gene conversion behaves similarly to selfing in that *t*_*b*_ is half the standard value (*N* versus 2*N*) and that *t*_*w*_ is much smaller than *t*_*b*_. This reduction in coalescence time with gene conversion can occur at much larger rates of sex relative to population size (*σ* ≫ 1*/N*) than the increase in coalescence time that we discussed earlier, though the absolute rate of sex must be low (*σ* ≪ 1).

The reason that gene conversion is so important when sex is low can be understood by considering coalescence from two samples taken from separate individuals. In the absence of gene conversion, two samples are put into same individual, on average, two separate times (requiring at least one bout of sex) before coalescing (e.g. the first time the two samples descended from different homologous chromosomes but the second time they descended from the same one). When the rate of sex is low, each time the two samples are put into the same individual, they persist together in that genotype for many asexual generations. If gene conversion is high relative to the rate of sex, the samples are likely to coalesce via gene conversion before the genotype is broken apart by sex. Thus, two samples need only to be put in same individual once before coalescing, so the coalescence time is half what it would be otherwise. If two samples start in the same individual, then the time to coalescence is simply the waiting time until gene conversion, 1*/γ* (which we have assumed is negligible to the waiting time for two alleles in separate individuals to enter the same ancestral individual, i.e., *N* generations). Note that when gene conversion is rare relative to sex (*γ →* 0), Equation 9 collapses to the results assuming no gene conversion (Equation 4). The effect of gene conversion on coalescent time is shown in Figure 3, demonstrating how it greatly reduces the coalescent time when it is strong compared to sex.

**Figure 3:**
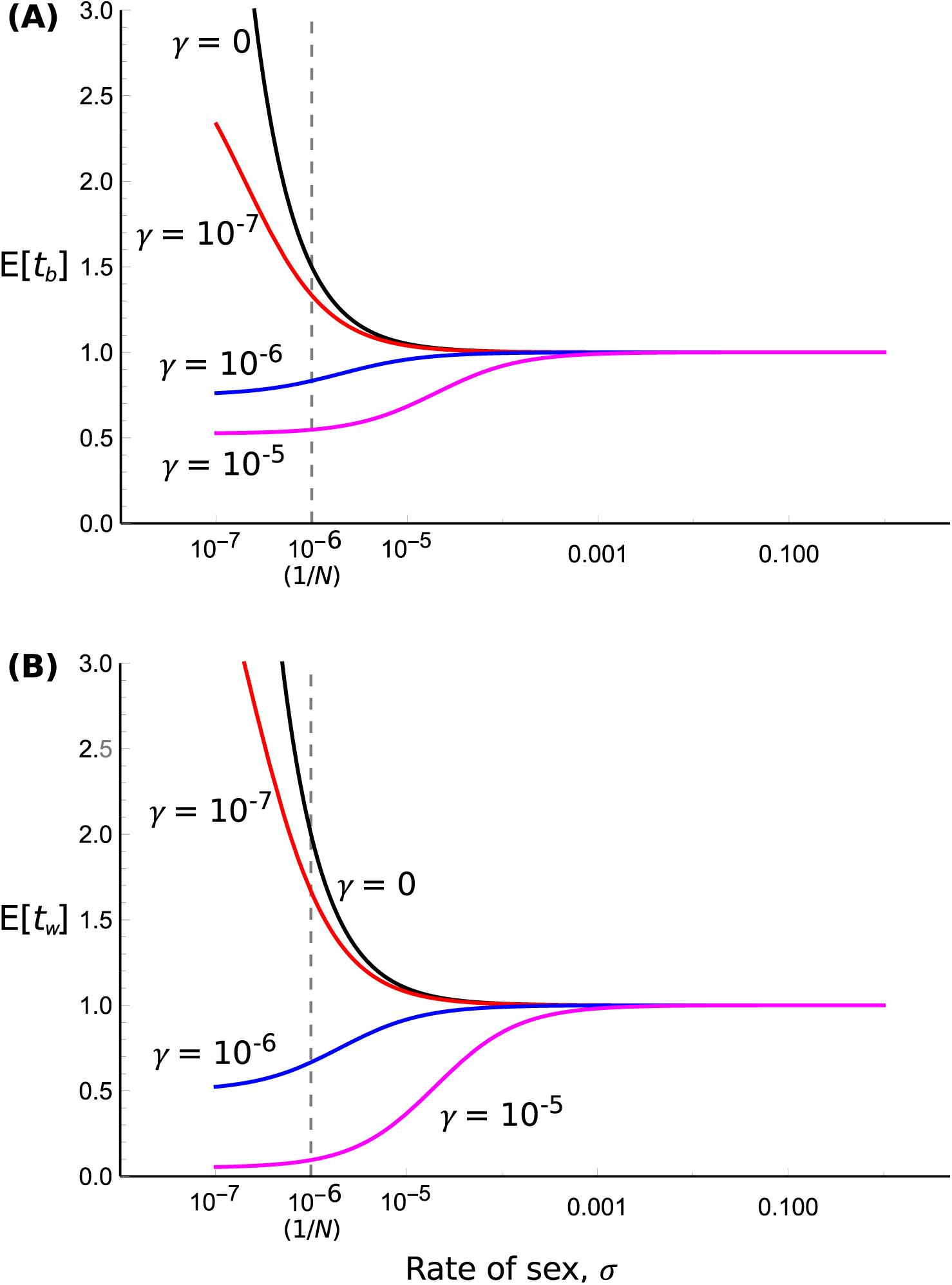
Expected coalescent time, relative to 2*N* generations, as a function of the rate of sexual reproduction. The population is of size *N* = 10^6^ for two samples taken from separate individuals (a) or the same individual (b). Gene conversion rates of *γ* = 0 (black), 10^-7^ (red), 10^-6^ (blue), and 10^-5^ (magenta) are shown.

Gene conversion rates are generally weak, with Flot *et al.* (2013) estimating gene conversion rates of 10^-5^ to 10^-6^ per site in bdelloid rotifers. However, these rates are feasibly the same order (or higher) as 1*/N*_*e*_ for most species, which the same rate at which infrequent sex starts affecting coalescence times. Thus the strong impact of gene conversion when sex is rare can explain why little within-individual divergence has been reported in facultative sexual genomes, despite very low (or zero) rates of sex (Crease and LYNCH 1991; Sch Ö N *et al.* 1998; Normark 1999; SchÖN and Martens 2003; Flot *et al.* 2013).

In multisample simulations, we found that low rates of gene conversion (*γ* = 1*/N*) generally lead to the opposite behaviour of summary statistics shown in Figure 2, due to the increased homozygosity. That is, most estimators decrease with lower sex, except for Fay and Wu’s H. The number of unique haplotypes also decreases for low rates of sex, as in cases without gene conversion (Figure S2 of the Supplementary Derivations file). Therefore the simultaneous reduction in both haplotype and genotype diversity with decreased rates of sex, in contrast to observing allelic sequence divergence, can be used to determine the presence of gene conversion (or a similar effect such as selfing) in facultative sexuals.

## Coalescent times for an island model

Here we show how a simple demographic model, the island model, can also be implemented into the facultative sexual analysis. The population consists of *d* demes, with *N*_*d*_ in each, so the total population size is *N*_*T*_ = *d × N*_*d*_. Each generation, an individual can migrate to another deme with probability *m* = *O*(1*/N*_*d*_) *≪* 1; this assumption ensures no more than one sample migrates per generation. To keep solutions tractable, we do not consider selfing in this analysis.

In this model, two alleles can be found in three non-coalesced states. Pairs of samples can either be taken from different demes; from different individuals within a deme; or from the same individual. We describe in Section S5 of the Supplementary Derivations file how the transition matrix **G** is formed, and solved to obtain coalescent times for different demes *t*_*d*_, within-deme *t*_*b*_, and within-individual *t*_*w*_. Conveniently, in the limit of low sex, migration, and gene conversion (1*/N ≪ σ*, *γ*, *m ≪* 1), *t*_*b*_ and *t*_*w*_ are exactly as in Equation 9 with *N* = *N*_*d*_*d*, and *t*_*w*_ is the same as *t*_*b*_ but with an additional (*d -* 1)*/*2*m* term to denote isolation by distance (Slatkin (1991); see also Section S2 of the Supplementary Mathematica file). Hence the effects of population structure, and the joint effect of gene conversion and low sex, are simply additive. As in a non-subdivided population, sex has to be on the order of the total population size, 1*/N*_*T*_ = 1*/*(*N*_*d*_*d*) in order to have a significant impact on coalescent time. If gene conversion is present with rate *γ ≈* 1*/N*_*T*_, then coalescence times are greatly reduced. Hence even a small rate of gene conversion reduces within-individual diversity that would otherwise be caused by low rates of sex. Further information is available in Section S2 of the Supplementary Mathematica file.

For multi-sample genealogies without gene conversion, several distinct types of tree topologies can occur depending on the relative rates of sex and migration. A standard genealogy with obligate sex and high migration is shown in Figure 4(a). It is well known (Wakeley 2000; Wakeley and Aliacar 2001) that with obligate sex and limited migration, samples separate themselves into subclades principally by their geographic location (Figure 4(b)), meaning that most of the variation is among demes; the same result occurs with facultative sex provided that sex is not too rare. On the other hand, if sex is low, then there tends to be two deep lineages, present within all demes, representing the long coalescent times between left/right haplotypes, assuming gene conversion is rare relative to sex (Figure 4(c)). That is, most of the variation is within individuals created by excess heterozygosity. When migration and sex are both low but migration is high relative to sex, then principally two major subclades arise based on the chromosome arm of the sample. Within these major clades, five further clades are apparent due to the geographical location of the samples (Figure 4(d)). A large fraction of the genetic variance is within individuals but there is also a considerable amount of variation among populations. Finally, if migration rates are the same order, or lower than the rate of sex, the opposite pattern is seen: there are five clades due to geographical separation, with pairs of subclades then forming due to lack of genetic segregation (Figure 4(e)). The frequency spectra for each of these five examples are shown in Figure S3(a)–(e) in the Supplementary Derivations file. When sex is low, samples present in half the individuals are very common, representing ongoing divergence at the two different allele copies. Furthermore, once isolation by distance is additionally prevalent, polymorphisms tend to arise in multiples of five, reflecting both segregation by deme and by allelic copy.

**Figure 4:**
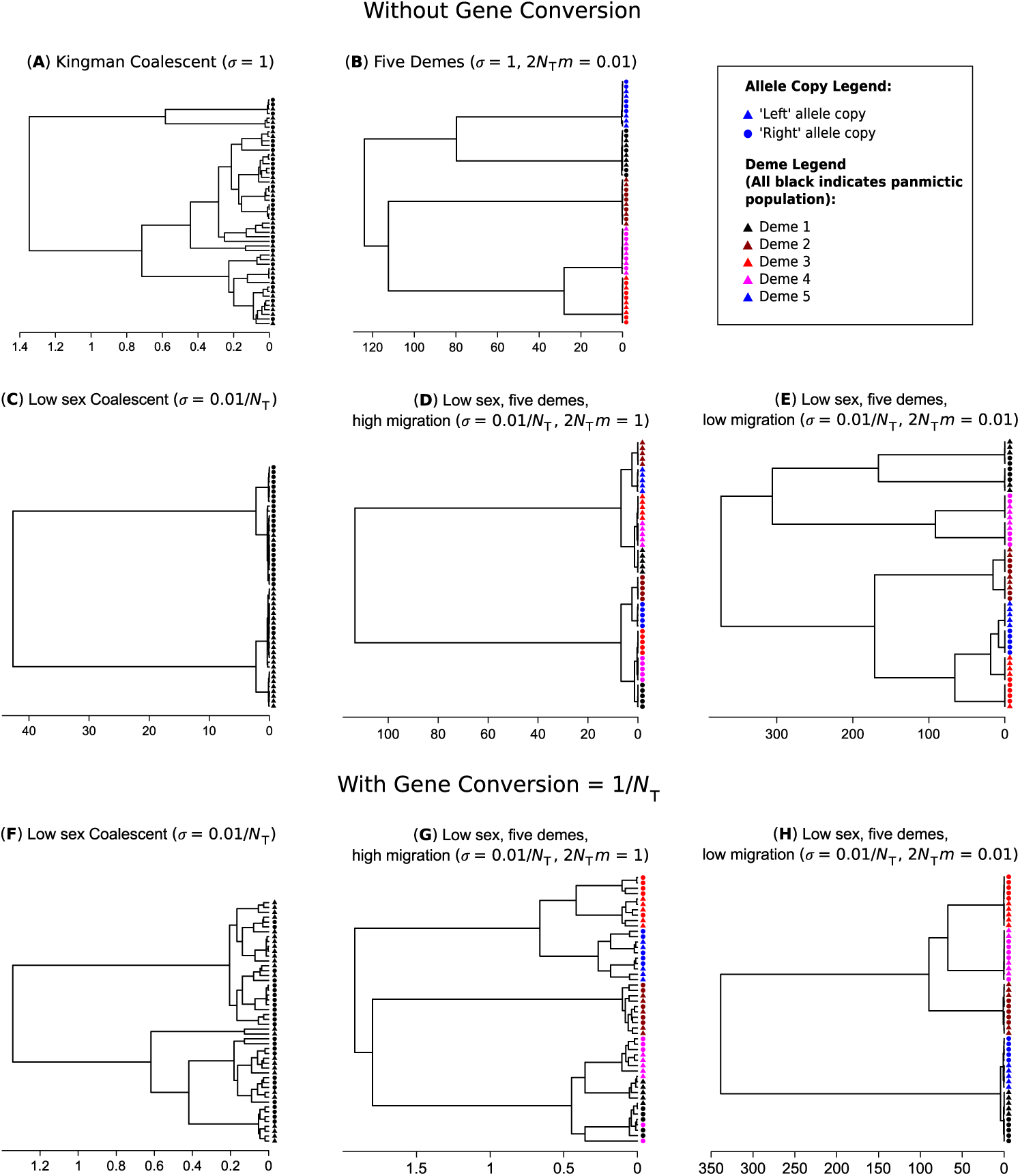
Examples of key genealogies produced by facultative-sex coalescent simulations. For all simulations, *N*_*T*_ = 10, 000 and 25 paired samples were simulated. Scale bars indicate time in past in units of 2*N*_*T*_ generations. Subdivision by either allelic copy or deme are shown in the legend. (a) Traditional (Kingman) coalescent, with a panmictic population and obligate sex. (b) Obligate sexual population in an island model, with *d* = 5 and 2*N*_*T*_ *m* = 0.01. (c) and (f) Panmictic population with low sex *N σ* = 0.01. (d) and (g) Low sex population (*N*_*T*_ *σ* = 0.01) in an island model with a relatively high migration rate (*d* = 5, 2*N*_*T*_ *m* = 1). (e) and (h) Low sex population (*N*_*T*_ *σ* = 0.01) in an island model with a relatively low migration rate (*d* = 5, 2*N*_*T*_ *m* = 0.01). No gene conversion is present for panels (a) to (e), but it is present at a rate *γ* = 1*/N*_*T*_ in panels (f) to (h).

With even small rates of gene conversion (*γ* = *O*(1*/N*_*T*_)), spatial structure remains but little genetic partitioning is evident, leading to greater mixing of the two chromosome arms (Figure 4(f)–(h)). Such trees give a misleading appearance of either a higher overall rate of sex, or spatially heterogeneous rates of sex. Figure S3(f)–(h) in the Supplementary Derivations file shows the relevant site frequency spectra: low-frequency variants are much more likely to be produced as in the obligate sex case, highlighting how even weak gene conversion creates spurious genomic signatures of frequent sex.

### Diversity measurements and parameter estimation

In the absence of gene conversion and with low rates of sex, it is possible to use the migration model results with *γ* = 0 to derive diversity-based estimators for the mutation, migration, and sex rates. In Section S5 of the Supplementary Derivations file, we outline how to derive the following scaled estimators for the mutation rate 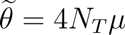, sex rate 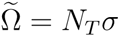, and migration rate 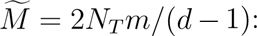:

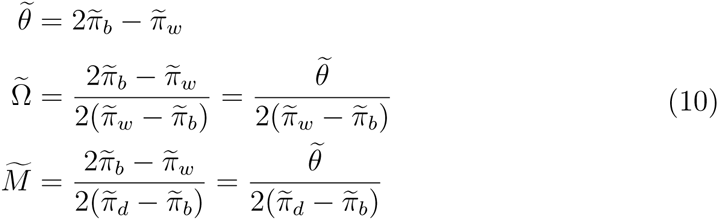

Here, 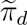 is the average between-deme pairwise diversity; 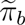 the within-deme diversity, and 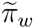 average within-individual diversity. We also investigate what sampling regimes should be used to most accurately estimate these parameters. Briefly, paired within-individual samples should be used as much as possible, and if taking the mean of Ω and *M* from multiple replicates, the ratio of means should be taken instead of the means of ratios. However, these estimators assume that within-individual coalescence times are the longest due to low rates of sex. If gene conversion occurs on the same order or greater than the rate of sex, within-individual diversity would be removed so these estimators perform badly. Hence such estimators would only work in the case where *γ ≪ σ ∼* O(1/*N*_*T*_), which may not apply to many partially asexual species that have large population sizes.

## Heterogeneity in rates of sex

Results so far have assumed that the rate of sex remains constant over genealogical time. However, this strong assumption may not be realistic for a broad range of taxa. Facultative sexual organisms are frequently observed to adjust their rate of sexual reproduction, depending on certain biotic and abiotic conditions that can vary in time and space. We outline how to extend the above derivations to account for either temporal or spatial heterogeneity. We do not consider gene conversion in these derivations.

### Temporally heterogeneous sexual reproduction

We consider a case where there is a single population, but the rates of sex change stochastically between two values over time. The same approach could be extended to allow any finite number of possible sex rates. Switches from periods when sex is at the lower rate *σ*_*L*_ to periods when it is at the higher rate *σ*_*H*_ occur at rate *p*_*LH*_; transitions in the reverse direction occur at rate *p*_*HL*_. To study coalescence with temporal heterogeneity, we assume that each of the two states in the original model can be found in low or high sex phenotype, resulting in four states.

We can also calculate the between-and within-individual coalescent times, as averaged over the two sex states, as follows:

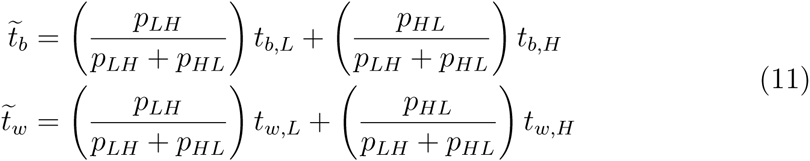

where *b*, *w* indicate between-and within-individual times, and the *L*, *H* subscripts further indicate coalescent times if in a low-sex or high-sex state. Calculating averages in this manner is only meaningful if switches between sex rates are probable relative to the coalescent time (i.e. *N p*_*LH*_, *N p*_*HL*_ *>* 0.1).

Solving to obtain the coalescent times yields solutions that are rather cumbersome (Section S3 of the Supplementary Mathematica file). However, we can show that over most parameter ranges, the coalescent times for the high-sex and low-sex regimes can be well-approximated by:

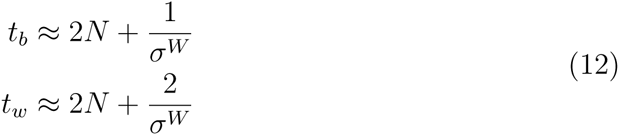

with *σ*^*W*^ is the weighted mean of the rate of sex dependent on how much time is spent in the high-sex and low-sex times:

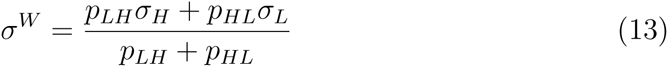

That is, these coalescent times are approximately equal to the constant-sex population (“non-temporal”) results (Equation 4), but with *σ* replaced by a weighted mean, depending on how much time is spent in the high-sex and low-sex states accordingly. We can show that such an approximation is directly obtained in the special case where both rates of sex are weak (*σ*_*L*_, *σ*_*H*_ = *O*(1*/N*)). Equivalent results arise if assuming that *σ*_*L*_ and *p*_*LH*_ = *O*(1*/N*), but no assumption is made on *σ*_*H*_; that is, where the population is generally asexual, but can undergo rare bursts bursts of obligate sex over short periods (Section S5 of the Supplementary Derivations file). Furthermore, if there are frequent transitions to obligate (or near obligate) sex, then coalescent times will be close to 2*N* generations in both the temporal and non-temporal cases. Trivially, calculating coalescent times using the weighted mean in this case provides a good approximation, since all solutions will be close to the obligate sex result of 2*N* generations. The main case where Equation 12 breaks down is if the transition probabilities *p*_*LH*_, *p*_*HL*_ are very small. In such a scenario, coalescence is likely to have occurred before any transition in sex rates arise, so Equation 12 would inaccurately reflect coalescence time. Figure 5 shows contour plots of the ratio of the exact values to the weighted mean approximation, verifying the parameter space as to when using the panmictic population result using the weighted mean of rates captures the coalescent time.

**Figure 5:**
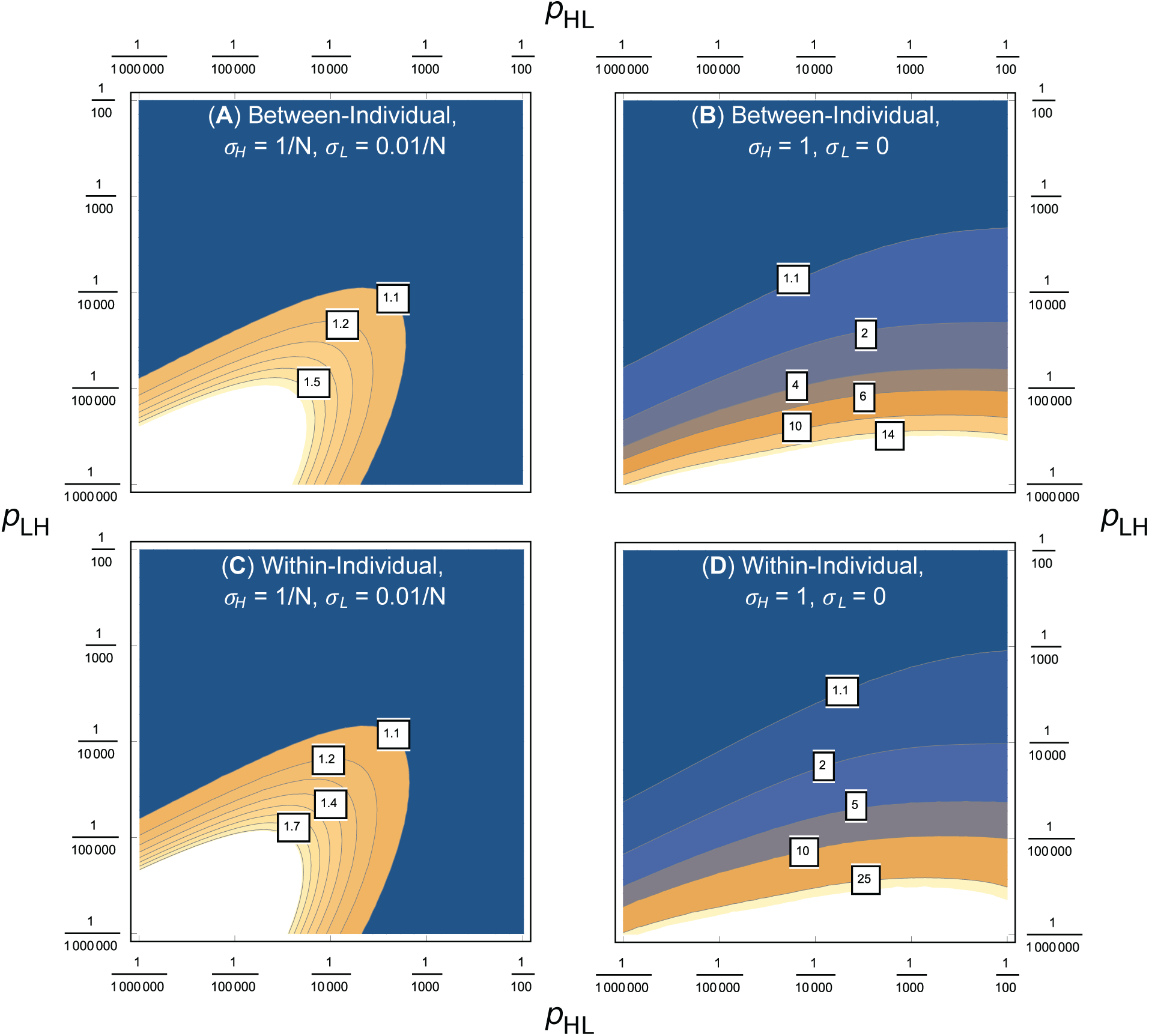
Contour plots of the ratio of exact coalescent times in a temporally heterogeneous population (Equation 11), to those predicted by Equation 12 (that is, using standard coalescent times with a weighted mean of sex rates). A value much greater than one indicates that the weighted mean solution greatly underestimates the true coalescent time. Plots are presented as functions of the transition probabilities *p*_*HL*_ and *p*_*LH*_. Some contour labels are added for clarity. (a) and (b) are for the between-individual case, with (c) and (d) the within-individual case. Other parameters are *N* = 10, 000, and *σ*_*H*_ = 1*/N*, *σ*_*L*_ = 0.01*/N* in (a) and (c), or *σ*_*H*_ = 1, *σ*_*L*_ = 0 in (b) and (d).

#### Analysis of individual coalescent times

If temporal heterogeneity is present, genome samples are usually taken from just one time period, and it is generally unknown when and how the rates of sex change. Based on this fact, we investigated how individual coalescent times (i.e. from either a low-sex or high-sex state, without averaging) differ in comparison to the non-temporal results. We show in the Section S3 of the Supplementary Mathematica file that, assuming *p*_*LH*_, *p*_*HL*_ *≪* 1, the four coalescent times can be written relative to the non-temporal results as follows:

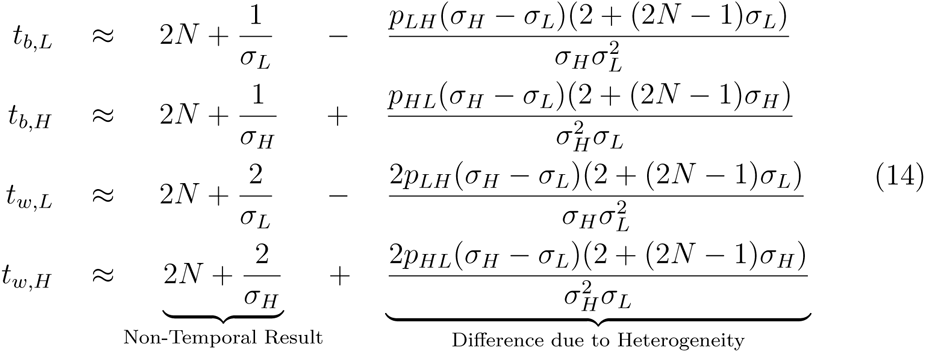

Three main results are apparent from investigating the differences between Equation 14 and the non-temporal solutions (Equation 4). First, low-sex differences are always negative, and high-sex differences are always positive. Hence temporal heterogeneity will decrease and increase the low-sex and high-sex times respectively, relative to the value expected based on the rate of sex at the time the alleles were sampled (the “non-temporal” case). Second, these differences are proportional to the difference between low and high-sex rates. Third, the additional terms depends only on the transition probability away from the current level of sex (for example, *t*_*b,L*_ is dependent on *p*_*LH*_ but not *p*_*HL*_). Overall, if one suspects that temporal heterogeneity affects the organism under study, then Equation 14 can be used to predict the actual coalescent times, given that genealogies will generally average over any heterogeneity.

In Section S5 of the Supplementary Derivations file, we apply estimators for the rate of sex (Equation 10) to simulation data obtained from a temporally heterogeneous population. If the rate of sex changes slowly over time, or if it shifts in a step-wise manner, then the average rate of sex will not be measured correctly. Conversely, if both rates of sex are small (*N σ* = *O*(1) or less), or the population switches often enough to the obligate sex rate (e.g. with *p*_*HL*_ = *p*_*LH*_ = 0.02), then estimates of Ω appears to reflect the weighted mean value.

### Spatially heterogeneous sexual reproduction

We next consider the case where there is an island model, but rates of sex occur at a lower rate *σ*_*L*_ in some demes and a higher rate *σ*_*H*_ in others (and these rates stay constant within each deme over time). There are *d*_*L*_ low-sex demes and *d*_*H*_ high-sex demes, with *d*_*L*_ + *d*_*H*_ = *d*_*T*_. Though we have assumed only two rates of sex, the same approach can be extended to consider an arbitrary number of rates.

When calculating pairwise coalescence times, there are now seven states to consider. For between-deme comparisons, two samples can either be taken from two different low-sex demes; two different high-sex demes; or one from a low-sex deme and the other from a high-sex deme. In addition, for within-deme samples (either from the same or separate individuals), they can either be from a high-sex or a low-sex deme.

One can form the 7 *×* 7 **G** matrix, but in this case deriving each term is complicated, and the overall solutions are huge. Hence the full derivation is saved for Section S4 of the Supplementary Mathematica File. However, important in-sights can be gained from the special case assuming that migration *m ≪* 1 and *σ*_*L*_ *≪ σ*_*H*_ *≪* 1. Here, the between-deme coalescent times for two high-sex demes (*t*_*D,HH*_), a high-sex and low-sex deme (*t*_*D,HL*_), and two low-sex demes (*t*_*D,LL*_) are obtained as:

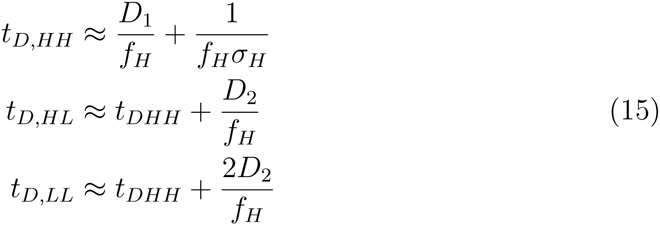

where *f*_*H*_ is the fraction of high-sex demes *d*_*H*_ */d*_*T*_, and *D*_1_, *D*_2_ are compound parameters:

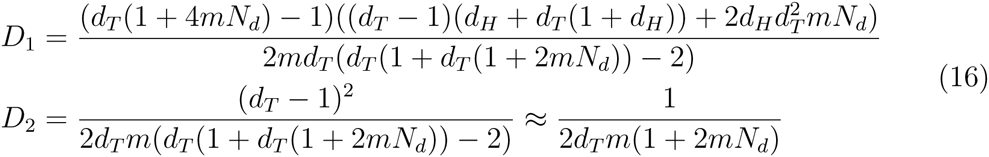

The approximation for *D*_2_ arises as *d*_*T*_ becomes large. Similarly, the approximate coalescent times between-individuals in a high- or low-sex deme (*t*_*b,H*_ and *t*_*b,L*_ respectively), and within-individual times (*t*_*w,H*_ and *t*_*w,L*_) can be built up as:

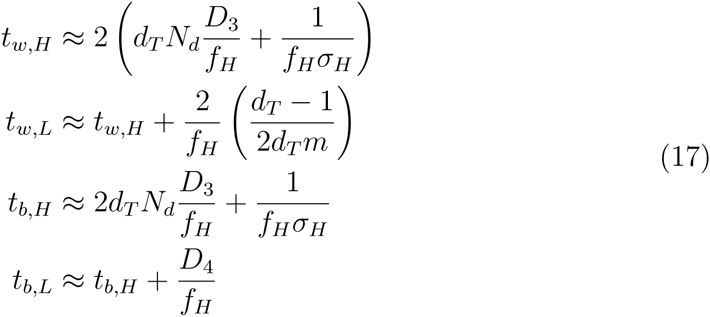

with *D*_3_, *D*_4_ being further compound parameters:

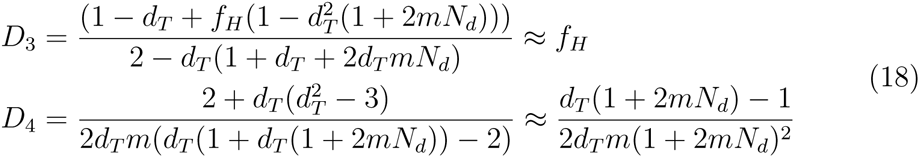

Again, approximations are for large *d*_*T*_. Furthermore, due to the recursive nature of Equation 17, elegant terms can also be obtained for the difference in coalescent times between high-sex and low-sex demes:

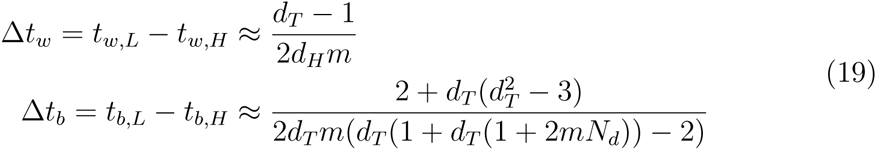

Equation 19 illustrates two points regarding how the difference in coalescent times for samples from low-sex demes compared to those from high-sex demes. The differences increase as migration rates decrease, and it is also independent of the rates of sex. This is because we have assumed *σ*_*L*_ *≪ σ*_*H*_ *≪* 1. Consequently, alleles from different left/right haplotypes cannot, or are extremely unlikely to coalesce within the low sex demes. Such alleles must first move to high sex demes before coalescence can occur. The difference shown in Equation 19 represents the waiting time for alleles in low-sex demes to migrate to high-sex demes before coalescence can occur. Equation 19 reflects the true differences, as long as *σ*_*L*_ is low enough compared to the migration rate (Section S4 of the Supplementary Mathematica file).

Another simple case we can consider is where there exists a single deme of each type (*d*_*L*_ = *d*_*H*_ = 1), and where both *σ*_*L*_ and *σ*_*H*_ are small but of the same order. Here, the difference in coalescent times for samples from low-sex demes compared to those from high-sex demes becomes:

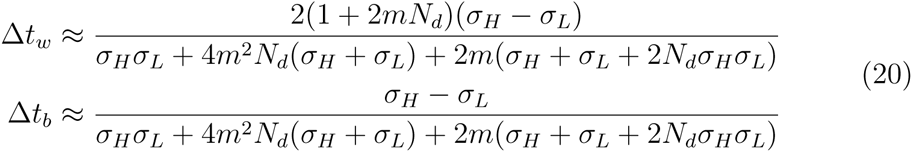

In this case, we can see that the difference in coalescence times grows with the difference in sex rates, and decreases with migration.

**Comparison with non-heterogeneous results.** We can ask how average coalescence times cc a metapopulation without spatial heterogeneity in sex rates, but with the same average rate of sex (i.e., using *σ* = (*d*_*L*_*/d*_*T*_) *σ*_*L*_ + (*d*_*H*_ */d*_*T*_) *σ*_*H*_). Figures 6(a) and 6(c) demonstrates that if the rate of migration and low-sex rates are not too low (2*N*_*T*_ *m* = 1 and *N*_*T*_ *σ*_*L*_ *≈* 1 respectively), then the average spatially heterogeneous within-individual coalescence time can be well-approximated by using a mean of sex rates. However, as both the migration rate and low-sex rates decrease, then using non-heterogeneous results with the mean rate of sex greatly underestimates the true coalescence time. Plotting the ratio of the coalescence times for low-and high-sex demes respectively demonstrate that coalescence times in the low-sex case increases disproportionally with reduced *σ*_*L*_ and *m*, compared to those in high-sex demes (Figures 6(b), 6(d)). Similar results arise for between-deme cases, as well as those for *σ*_*H*_ = 1 (Section S4 of the Supplementary Mathematica file).

**Figure 6:**
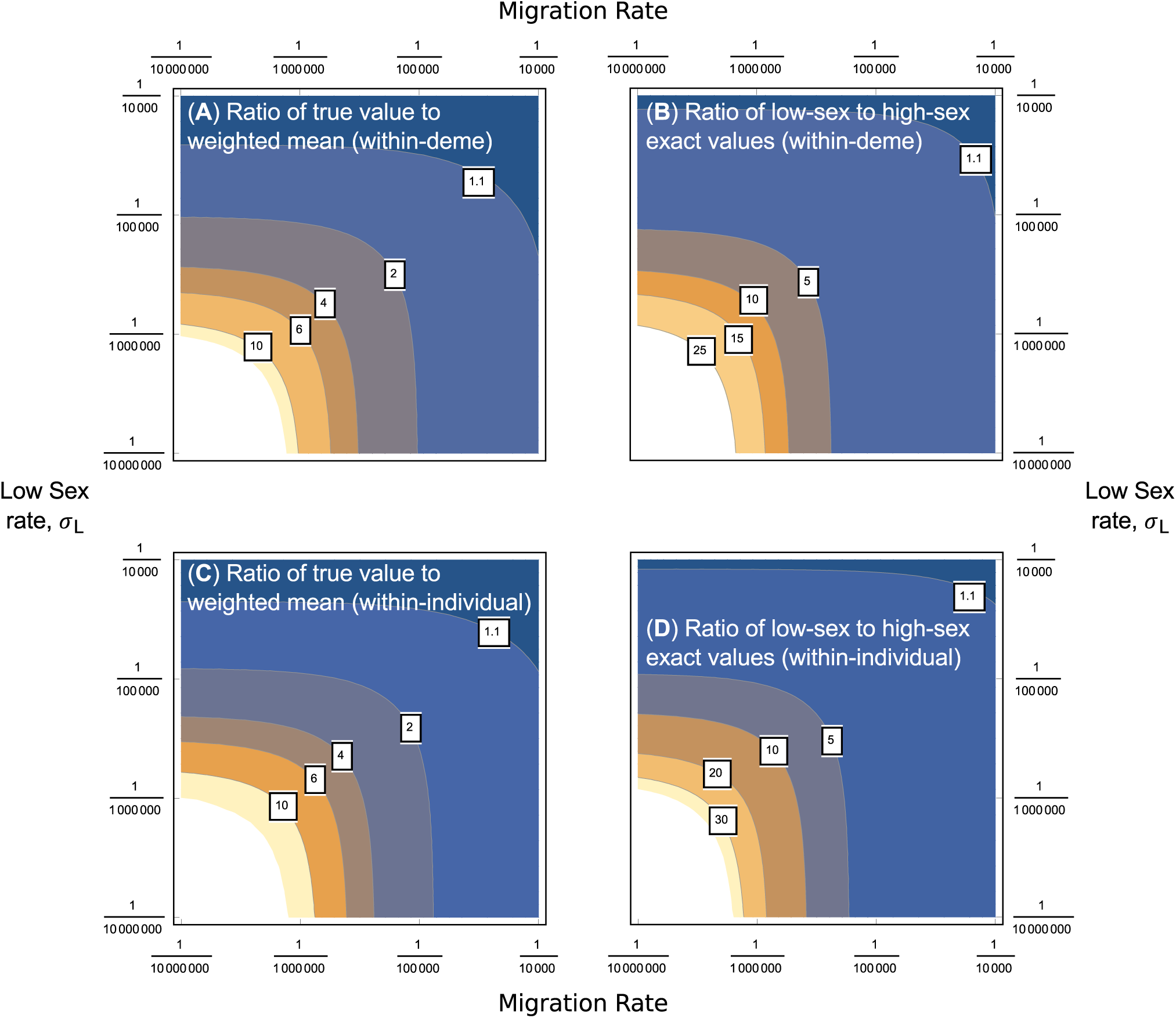
Contour plots of the ratio of either the exact within-individual coalescent times in a spatially heterogeneous population to those predicted by a weighted mean of sex rates ((a) and (c)), or the ratio of the exact coalescence times in low-sex demes compared to high-sex demes ((b) and (d)). Plots are presented as functions of the migration rate *m* and the low rate of sex *σ*_*L*_. Some contour labels are added for clarity. (a) and (b) are for the between-individual case, with (c) and (d) are the within-individual case. Other parameters are *N* = 10, 000, *σ*_*H*_ = 1*/N*, *d*_*T*_ = 4 with *d*_*H*_ = *d*_*L*_ = 2.

In Section S5 of the Supplementary Derivations file, we apply diversity-based estimators for the rates of sex to simulation data from a spatially heterogeneous case, in the absence of gene conversion. If migration is not too high (2*mN*_*T*_ *≪* 1), then application of these estimators to individual demes can give a decent estimation of the rate of sex within them. Estimates are less accurate if migration is higher (2*mN*_*T*_ *∼* 1), instead predicting a rate of sex intermediate between the two true values.

## Discussion

We have outlined here a comprehensive theoretical treatment to consider coalescent processes with arbitrary rates of sex. After re-applying structured coalescent theory to previous models (Bengtsson 2003; Ceplitis 2003), we show how to calculate expected coalescent times for pairs of samples, either taken between-or within-individuals (Equation 4). We show how this structured coalescent can be further extended to take other cases into account, such as self-fertilisation (Equation 6), gene conversion (Equation 8), and migration according to an island model. From the migration case, we further derived statistical estimators for the rates of mutation, migration, and sex (Equation 10).

Our multi-sample simulation results highlight that, with very low rates of sexual reproduction, strong departures from the standard neutral model expectations for summary statistics such as Tajima’s D and Fay and Wu’s H are expected, reflecting the deep genealogical divergence between alleles within individuals. Thus, species with very low rates of sex can exhibit diversity patterns resembling long-term balancing selection and/or population subdivision. This highlights the importance of analysis of within versus between individual patterns of diversity, to help untangle the relative role of demographic history and selection from partial sex on the shape of genealogies.

To more accurately account for demographic effects, we found two solutions. First, newly derived estimators for the mutation, sex, and migration rates based on pairwise diversity (Equation 10) can be accurate, if good sampling practice is followed (Section S5 of the Supplementary Derivations file). However, these simple estimators can produce misleading results under a wide variety of scenarios that cannot be accounted for, due to insufficient degrees of freedom in the equations. These include the presence of gene conversion rates of similar magnitude to sex, and spatial or temporal heterogeneity in sex. We therefore suspect that the best manner to account for complex demography would be to use coalescent simulations in an inference framework, such as Approximate Bayesian Computation (Abc; SunnÅKer *et al.* (2013)), to predict how summary statistics change under different regimes. Our new simulation package allows for ABC to be performed while varying the rate of sex, to simultaneously infer sex and demography.

One major result is that once sex becomes on the same order as gene conversion, the latter becomes a powerful force in reducing within-individual diversity, and therefore removing genetic signatures of rare sex. Beyond the challenges this presents for estimating rates of sexual reproduction, this result also has important evolutionary consequences since the absence of permanent heterozygosity can influence not only the structuring of neutral diversity but also the exposure of both deleterious and advantageous recessive mutations to selection. Although elevated heterozygosity has been observed in some partial and obligate asexual organisms (Tucker *et al.* 2013; Hollister *et al.* 2015), there is increasing evidence that a major source of high within-individual diversity may be hybrid origins of asexual lineages, rather than ongoing accumulation of diversity due to new mutations (Tucker *et al.* 2013). Under biologically realistic rates of gene conversion, this initially high heterozygosity may be rapidly eroded by gene conversion, decreasing neutral variation within and between individuals (Tucker *et al.* 2013).

Since it is also known that many organisms exhibit plastic rates of sex, we have also included both spatial and temporal heterogeneity into our results. With temporal heterogeneity, the expected coalescent time can be well-approximated in many cases by using the constant-sex population result with a weighted mean rate of sex (Equation 12). Such an approach is invalid, however, if long periods of time are spent in a low-sex regime before transitioning arises. In this case, the approximations in Equation 14 can be used to estimate how coalescent times are extended or reduced in the presence of temporal heterogeneity. In addition, using a weighted mean approximation is generally invalid with spatial heterogeneity, since low-sex demes contribute disproportionally to coalescence (Figure 6). However, if migration rates are low and gene conversion is rare relative to sex, one can accurately measure rates of sex within each deme (Table S6 in the Supplementary Derivations file).

These analyses provide a more thorough theoretical basis for which the genealogies of samples from facultative sexuals can be reconstructed, in order to provide the null distribution of neutral diversity for such populations. We have outlined methodology so that it is clear how the results presented here can be extended to cover even more complex scenarios; nevertheless, there are several apparent routes for extending this work. Although we have showed how to implement demography via an island model, it would be desirable for future models to consider a broader array of demographic scenarios. Real-world populations go through a bewildering array of changes, including extinction, recolonisation, population expansion and bottlenecking (Whitlock and Mccauley 1999; Veeramah and Hammer 2014). Advanced coalescent models for obligate sexuals have been developed to determine how complex demographic effects affect genealogies (Rosenberg and Nordborg 2002; Hudson 2002); it will certainly be worthwhile to implement similar demographic scenarios into facultative sex coalescent simulations, given the known impact of spatial effects on clonal species distribution (Arnaud-Haond *et al.* 2007).

The other major extension that will prove important is to implement recombination. In this case, the outputs will no longer be a genealogy, but a recombination graph (Griffiths 1981, 1991), although it would still be possible to create genealogies for unrecombined genomic segments. Recombination graphs can provide greater power at detecting demographic effects with fewer samples, although it can be more computationally demanding to run relevant analyses (Schiffels and Durbin 2014). Adding recombination will prove important for the facultative sexual coalescent, because while sex has to be very rare (*O*(1*/N*)) to affect the non-recombining genealogies presented here, differences in recombination graphs might become apparent with higher rates of sex. This is because it is expected that recombination correlates linearly with sex rate. A similar result was derived for partially-selfing organisms by Nordborg (2000). Genomic sequences of facultative sexuals can reveal evidence of rare recombination events, providing proof of cryptic sex (examples were shown by Grimsley *et al.* (2010) for the marine algae *Ostreococcus* spp., and Signorovitch *et al.* (2005) for Placozoa). In addition, ancestral graphs would be able to separate out areas of the genome affected by gene conversion, leading to much more refined measurements of how this force affects genetic architecture. In particular, gene conversion events in the absence of sex will lead to much higher rates of gene conversion relative to crossing over. This will have a strong effect on short-range versus long-range linkage disequilibrium along chromosomes, which may enable accurate estimators of the rate of sex even in the presence of significant rates of gene conversion. Recombination graphs of facultative sexuals can thereby greatly increase the power by which the effects of infrequent sex and demography can be disentangled.

## Acknowledgements

MH is supported by a Marie Curie International Outgoing Fellowship, grant number MC-IOF-622936 (project SEXSEL). This work was also supported by Discovery Grants (AFA & SIW) from the Natural Sciences and Engineering Research Council of Canada.

